# A predictive model of intercellular tension and cell-matrix mechanical interactions in a multicellular geometry

**DOI:** 10.1101/701037

**Authors:** Lewis E. Scott, Lauren A. Griggs, Vani Narayanan, Daniel E. Conway, Christopher A. Lemmon, Seth H. Weinberg

## Abstract

Epithelial cells form continuous sheets of cells that exist in tensional homeostasis. Homeostasis is maintained through cell-to-cell adhesions that distribute tension and balance forces between cells and their underlying matrix. Disruption of tensional homeostasis can lead to Epithelial-Mesenchymal Transition (EMT), which is a transdifferentiation process in which epithelial cells adopt a mesenchymal phenotype, where cell-cell adhesion is lost and individual cell migration is acquired. This process is critical during embryogenesis and wound healing, but is also dysregulated in many disease states. To further understand the role of intercellular tension in spatial patterning of epithelial cell monolayers, we developed a multicellular computational model of cell-cell and cell-substrate forces. This work builds on a hybrid Cellular Potts-finite element model to evaluate cell-matrix mechanical feedback of an adherent multicellular cluster. Thermodynamically-constrained cells migrate by generating traction forces on a finite element substrate to minimize the total energy of the system. Junctional forces at cell-cell contacts balance these traction forces, thereby producing a mechanically stable epithelial monolayer. Simulations were compared to *in vitro* experiments using fluorescence-based junction force sensors in clusters of cells undergoing EMT. Results indicate that the multicellular CPM model can reproduce many aspects of EMT, including epithelial monolayer formation dynamics, changes in cell geometry, and spatial patterning of cell geometry and cell-cell forces in an epithelial colony.

**Author summary:** Epithelial cells line all organs of the human body and act as a protective barrier by forming a continuous sheet. These cells exert force on both their neighboring cells as well as the underlying extracellular matrix, which is a network of proteins that creates the structure of tissues. Here we develop a model that encompasses both cell-cell forces and cell-matrix forces in an epithelial cell sheet. The model accounts for cell migration and proliferation, and regulates how cell-cell adhesions are formed. We demonstrate how the interplay between cell-cell forces and cell-matrix forces can regulate the formation of the epithelial cell sheet, the organization of cells within the sheet, and the pattern of cell geometries and cell forces within the sheet. We compare computational results with experiments in which epithelial cell sheets are disrupted and cell-cell junction forces are measured, and demonstrate that the model captures many aspects of epithelial cell dynamics observed experimentally.

## Introduction

The epithelium is characterized by polarized sheets of cells that form by self-organization and reside in a mechanical equilibrium (reviewed in [1]). This mechanical equilibrium is maintained by regulation of both adhesion between neighboring epithelial cells (cell-cell) as well as adhesion between epithelial cells and the underlying extracellular matrix (cell-matrix). Cells generate cytoskeletal tension via actomyosin contractility, which is transmitted to the underlying matrix, while cell-cell adhesion mechanically couples abutted cells and distributes cytoskeletal tension to neighboring cells. This physical cellular interconnectivity and balance of tension at the cell-matrix and cell-cell interfaces produces a coupled monolayer that acts as a cohesive structure in static equilibrium.

Maintenance of static equilibrium in the epithelial sheet is essential to maintaining barrier and signaling functions of the epithelial sheet; however, disruption of the static equilibrium plays an important role in both physiological phenomena such as embryogenesis and pathological states including fibrosis and tumorigenesis [2, 3]. Mechanical equilibrium relies on tissue scale coordination of mechanical dynamics extending beyond local cell-cell and cell-matrix adhesions [4]. Local perturbations to the equilibrium state result in localized tension in the monolayer and a disruption to the equilibrium. For example, the cellular phenomena known as epithelial-mesenchymal transition (EMT), which is essential for embryogenesis and tissue morphogenesis, but which has also been implicated in tumorigenesis and fibrotic diseases, is initialized by perturbations in cell-cell adhesion. This process results in a phenotypic switch in which epithelial cells transdifferentiate into mesenchymal cells (reviewed in [5]). The perturbation in cell-cell adhesion redistributes tension in the monolayer, and cell-matrix adhesion compensates for the resulting localized stress. As such, spatial patterning of mechanical stress can facilitate phenotypic regulation and is crucial to both maintenance and disruption of tissue homeostasis [2, 4, 6].

Previous studies have explored the role of cell-cell adhesion in maintaining tensional homeostasis in the epithelial monolayer: increasing cellular contractility has been shown to stimulate formation of cell-cell junctions [7], and subsequent transfer of force to the cell-cell adhesion allows for stress distribution about the monolayer to maintain tensional homeostasis [4, 6]. As a result, mechanical gradients form that define spatial patterns and provide positional information within the monolayer. Both *in vitro* and *in silico* studies have demonstrated that the forces of a monolayer correspond to its geometry [8, 9].

In this work, we explore the role of cellular adhesion in maintaining tensional homeostasis of epithelial monolayers. To simulate epithelial monolayers, we extended a model developed by van Oers et al, which consists of a hybrid Cellular Potts model (CPM) and finite element model (FEM) [10]. The model simulates individual cellular traction forces based on their geometric size and shape, as has previously been modeled and validated by one of the senior authors of this work [11]: cellular traction forces are proportional to the first moment of area (FMA) about each point in the individual cellular geometry. This results in a pattern of traction forces directed towards the cell centroid and proportional to their distance from the cell centroid. These traction forces generate substrate strains which, in addition to cell-cell and cell-matrix interactions, impose a thermodynamic constraint and govern the dynamics of individual cells in the CPM. In the current work, we incorporate the formation of cell-cell adhesion between neighboring cells to accurately represent the biology of epithelial cells. We extend the Lemmon and Romer FMA model to multicellular clusters, and model traction forces based on the multicellular geometry rather than the individual cell. Thus, individual cell traction forces are proportional in magnitude to the distance from the centroid of the multicellular cluster, instead of the centroid of the individual cell.

In the original Lemmon and Romer model, each cell is in static equilibrium: because traction forces are proportional to the first moment of area, and the centroid by definition is the point where the integral of the first moment of area is zero, all traction forces within a cell must sum to zero. However, when we calculate traction forces based on the multicellular cluster, each individual cell is no longer in static equilibrium. Previous studies have suggested that cells in epithelial monolayers exist in a quasi-equilibrium, even when cell-cell junctional forces are present [7]. As such, we model the force applied to the cell-cell junction as the balancing force that opposes the traction forces for that cell, resulting in a quasi-equilibrium for each cell. This assumption has been observed experimentally in epithelial cell pairs, in which the junction force is equal and opposite to the net traction force [7]. We thus are able to predict the formation of an epithelial monolayer, including epithelial cell geometry, cell-matrix traction forces, and cell-cell junctional forces, based on first principles of cell contractility, cell geometry, and thermodynamic energy minimization. Results are compared to *in vitro* experiments in which epithelial monolayers were grown in a predetermined geometry established by microcontact-printed islands. Cell geometry and cell-cell junctional forces are quantified and compared to simulations. To further probe the role of junctional forces in tissue homeostasis, we induce phenotypic changes in epithelial clusters via addition of Transforming Growth Factor-*β*1 (TGF-*β*1), a known inducer of EMT. To replicate these effects in the model, we change the relative weight of cell-cell and cell-matrix interfacial energies in the CPM equations, and predict how changing phenotype can facilitate disruption of mechanics and morphology in the epithelial sheet.

Simulations demonstrate that traction forces of multicellular colonies scale linearly with the size of the colony, independent of the individual cell geometry. Additionally, we demonstrate that the model can be generalized to predict the distribution of junctional forces across a monolayer: junction forces are predicted by a quadratic function that is highest at the colony center and decays towards the cell edge. These predictions are independent of indvidual cell geometry and are consistent with existing literature [12].

## Results

### Multicellular traction forces drive formation of epithelial monolayers

Prior studies from van Oers and colleagues demonstrated that a hybrid CPM-FEM model can predict cellular spreading and organization based on cell-generated traction forces, resulting strains in the substrate, and durotaxis-driven migration in the CPM. To expand this model to adherent cell monolayers, we incorporated several advancements: first, cellular traction forces were predicted from the FMA model [11] based on a cell cluster geometry, not on individual cells. As such, cells in contact with neighboring cells “adhere” and begin to generate traction forces as a cohesive unit. Second, we assume that each cell in a multicellular cluster still maintains a static equilibrium, as has been suggested previously [7]. As such, we require the force acting on cell-cell junctions to counter the net traction force for each cell, as illustrated in a simple two cell example (Fig 1C, left).

**Fig 1.**
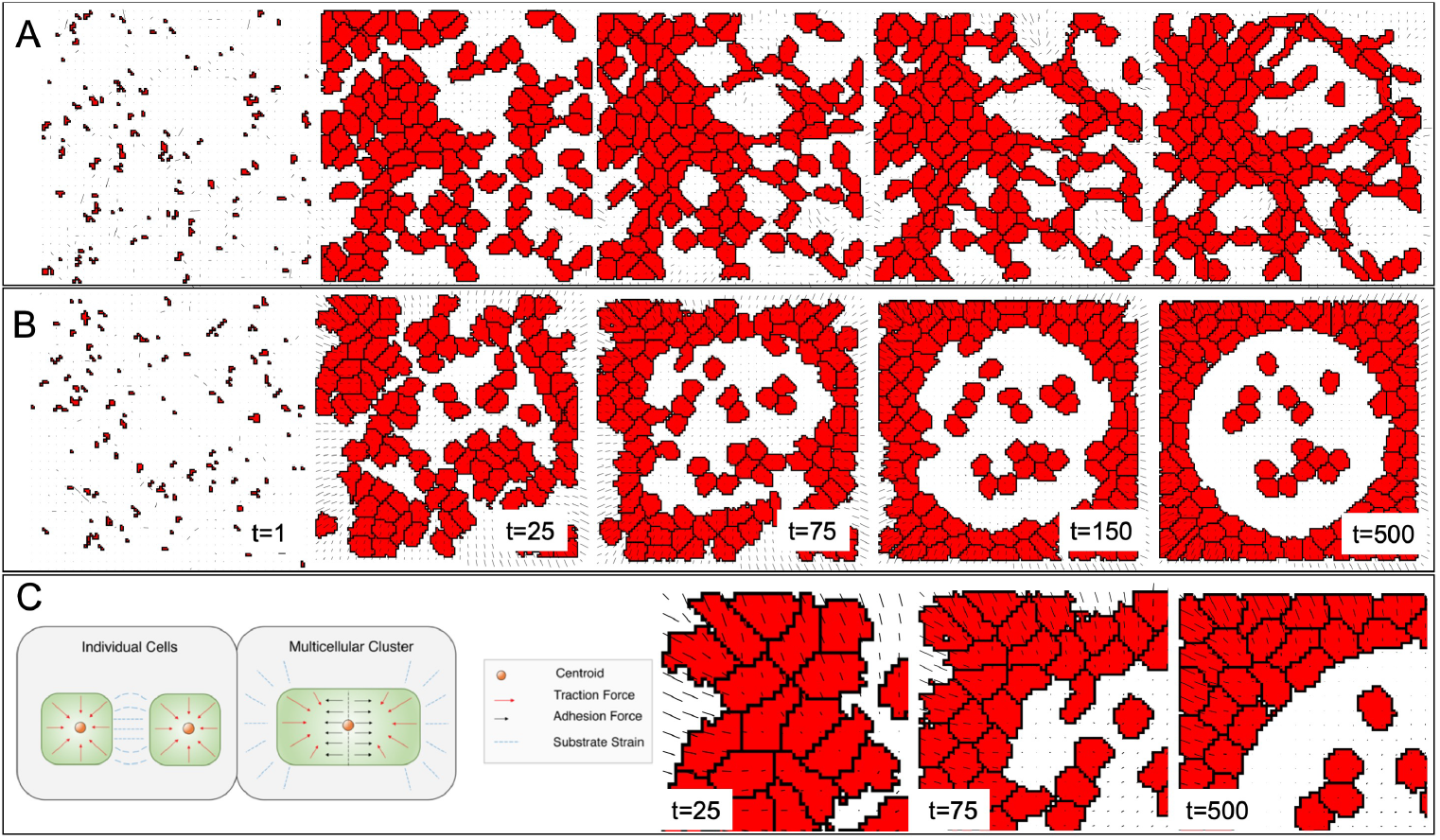
Simulated cells (red pixels) migrate on a finite element substrate that responds to cell-generated traction forces. Traction forces are calculated based on either (A) individual cell geometries or (B) multicellular clusters. (C, left) Representation of traction forces with resulting strain for multicellular geometries, and (C, right) inset of time points from panel B.

Figure 1 depicts simulated cells (red pixels) with corresponding scaled substrate strains (black vectors) for two scenarios. In the first, traction force is calculated from the first moment of area (FMA) about the single cell geometry and each cell is in static equilibrium. As a result, the net imbalance for each cell is zero and no force is transferred across the cell-cell junction (Fig 1A). In the second scenario, traction force is calculated from FMA about the multicellular geometry and each *cluster* is in static equilibrium (Fig 1B). The net force imbalance for each cell is balanced by the intercellular tension, which transfers the traction force to neighboring cells. Without redistribution of cytoskeletal stress to neighboring cells across cell-cell junctions, cellular alignment is localized and multicellular structures behave as partially cooperative networks with discordant substrate strains (Fig 1A, S1 Video), as demonstrated by van Oers et al [10]. In contrast, traction force distribution across cell-cell junctions to neighboring cells results in highly cooperative networks with a uniform spatial gradient of substrate strains. The formation of these cohesive multicellular clusters resembles an epithelial monolayer with preferential localization towards the boundary (Fig 1B, S2 Video). In the resulting multicellular clusters, net traction forces have a magnitude and direction at any given point proportional to the FMA about that point in the cluster, resulting in a linear gradient of substrate strain oriented radially towards the cluster centroid (Fig 1C, right, S1 Fig).

### Spatiotemporal dynamics of monolayer confluence

Preliminary simulations demonstrated the formation of a subconfluent monolayer-like sheet, which alters the spatial distribution of monolayer stress. To better predict the spatiotemporal dynamics of an *in vitro* epithelial monolayer, we incorporated cellular proliferation into the CPM to account for cell division dynamics, and then compared the spatiotemporal dynamics with cultured epithelial cells (Fig 2, S2 Video; see Methods for a more in-depth discussion). Mammary breast epithelial cells (MCF10A) were seeded onto poly-dimethyl siloxane (PDMS) substrates with a 250 *µ*m x 250 *µ*m microcontact-printed area of laminin (Fig 2A). Epithelial monolayers reached confluence over approximately 24 hours. Simulated cells exhibit similar patterning representative of MCF10A confluence dynamics (Fig 2B). To estimate the rate of proliferation in the simulations, immunofluorescence images were analyzed at 0, 6, 12, 18, and 24 hours and quantified for confluence as a function of time (Fig 2C; S3 Video). The half maximal confluence for simulations and experiments indicate that 1 Monte Carlo steps (MCS) corresponds to approximately 4.8 minutes of experimental time (Fig 2B, C). The experimental time scale was used to estimate a simulated division probability of 0.5% per MCS. These results demonstrate that simulated spatiotemporal dynamics approximate cellular dynamics observed *in vitro* and agree with previous studies [13].

**Fig 2.**
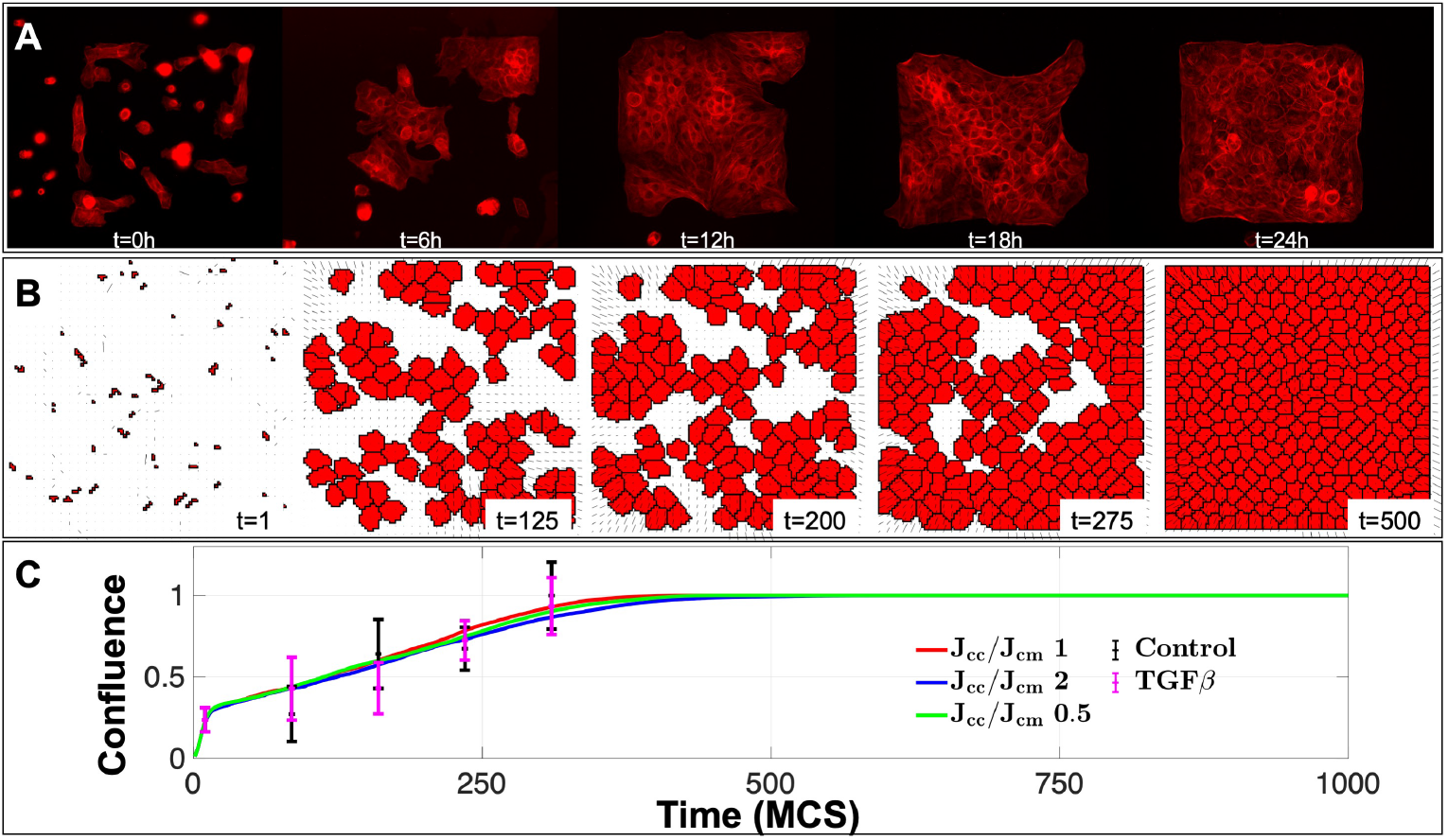
Spatiotemporal dynamics of simulated and *in vitro* tissue patterning. Visual comparison of time points from initial seeding to confluence illustrates parallels between (A) *in vitro* and (B) simulated spatial patterns. (C) Confluence, the fraction of total cell area to total substrate area, is shown as a function of time or Monte Carlo Steps (MCS), for *in vitro* and *in silico* experiments, for different conditions. Other parameters: Time scale: 4.8 min/1 MCS, *J*_*cm*_ = 2.5.

### Decreasing contact inhibition increases cell size and decreases cell number

With the key addition that traction forces are governed by the FMA model about the cluster geometry rather than the single cell geometry, the previous results illustrate distinct spatial patterning representative of epithelial monolayers. We next utilized our model to simulate epithelial monolayer and associated EMT-like dynamics. One key aspect of the epithelial phenotype is contact inhibition: that is, the propensity of a cell to stop migration when a neighboring cell is encountered [14, 15]. As epithelial cells undergo EMT and become more mesenchymal, contact inhibition is reduced [16]. To mimic the effects of EMT in epithelial monolayers in our multicellular FMA model, we varied the relative interaction energies between neighboring cells in the CPM, which simulates changes in contact inhibition. We varied the ratio of interaction energies at the cell-cell and cell-matrix interfaces, *J*_*cc*_ and *J*_*cm*_, respectively (see Materials and Methods, Eq 7), for the single cell (Fig 3A-D) and multicellular (Fig 3E-H) FMA models. The magnitude of the respective energies represents a prohibitive interaction, i.e., a higher *J*_*cc*_/*J*_*cm*_ ratio reflects increased contact inhibition between adjacent cells. For each simulation, we measured the steady-state monolayer confluence, average cell area, total cell count, and relative net cellular traction forces, averaged over 5 simulations with distinct random cell seeding, and plotted these measures as a function of the *J*_*cc*_/*J*_*cm*_ ratio. These simulations were then repeated for 3 distinct values of cell-matrix interaction energies, *J*_*cm*_.

**Fig 3.**
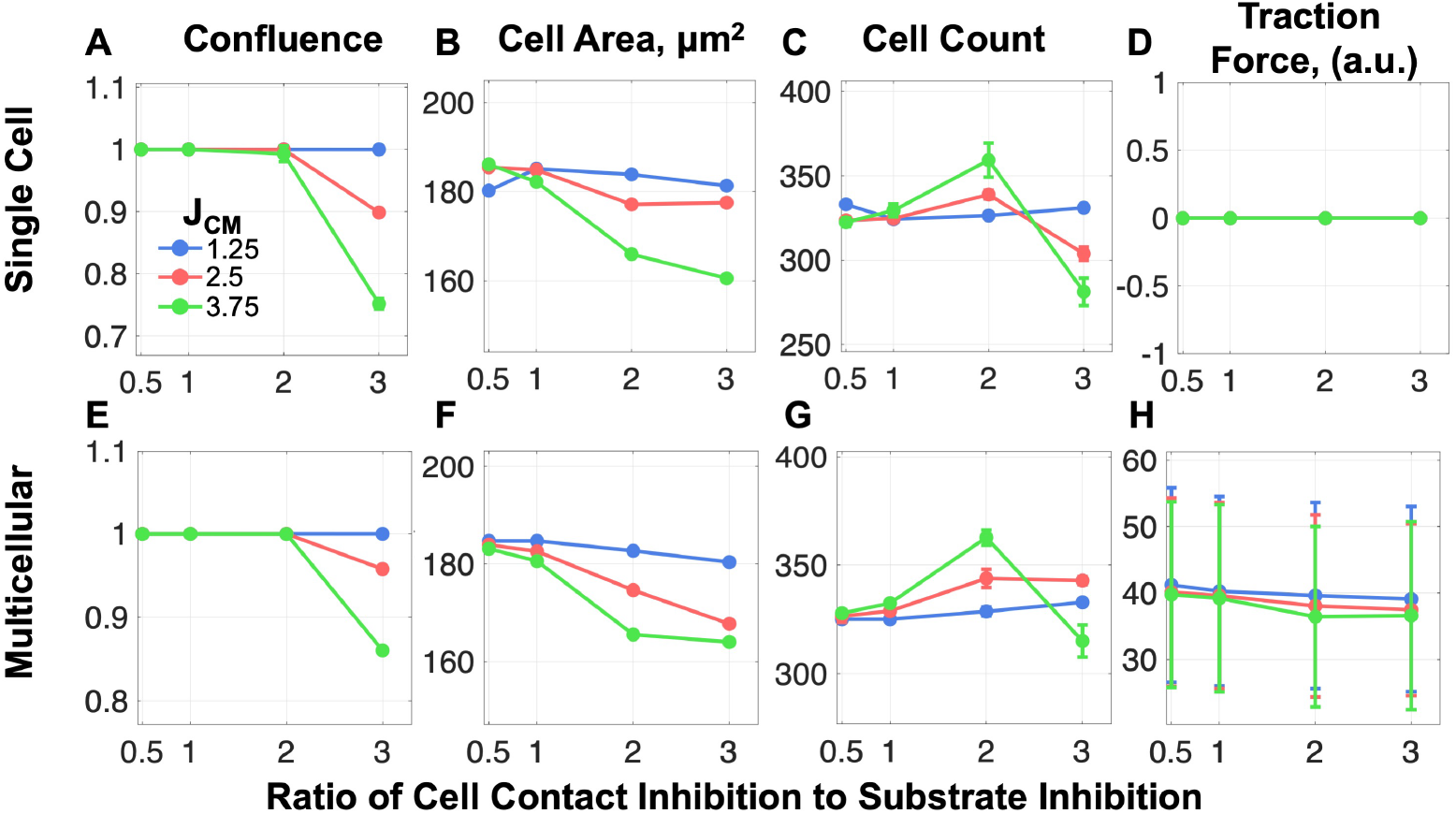
Parameter sweep of interaction energies. (A-D) Single cell FMA and (E-H) multicellular FMA simulated confluence, cell area, cell count, and traction force, shown as a function of the ratio of cell-cell contact inhibition to cell-matrix inhibition (*J*_*cc*_/*J*_*cm*_), varying *J*_*cm*_ values.

Results indicate similar trends between the single cell and multicellular FMA models, with the exception of net cellular traction force, which must be zero for a cell in static equilibrium in the single cell FMA model (Fig 3D). Beyond a critical point (*J*_*cc*_/*J*_*cm*_ = 2), high cell contact inhibition precludes the formation of confluent monolayers (Fig 3A, E). Further, we find that the time course of monolayer confluence only weakly depends on cell contact inhibition below this critical point, i.e. for conditions that form confluent monolayers (Fig 2C). Similarly, increasing cell contact inhibition results in smaller cell area (Fig 3B, F) and higher cell count (Fig 3C, G). In the multicellular FMA model, net traction force per cell decreases as the *J*_*cc*_/J_*cm*_ ratio increases. We find that higher substrate inhibition, i.e., increased J_*cm*_, tends to increase the sensitivity to the *J*_*cc*_/*J*_*cm*_ ratio for all measures. Thus, these data indicate that a loss of contact inhibition leads to larger cells, lower cell count, and in extreme cases, loss of confluence.

### Decreasing simulated contact inhibition mimics TGF-*β*1-induced EMT

The above results suggest that cells in the multicellular FMA model resemble the archetypal phenotype of epithelial cells undergoing EMT. With decreased cell-cell contact inhibition (i.e., smaller *J*_*cc*_/*J*_*cm*_ ratio), simulated cells exhibit the characteristic increased spreading and decreased proliferation of the mesenchymal phenotype, while at increased cell-cell contact inhibition (i.e., larger *J*_*cc*_/*J*_*cm*_ ratio), simulated cells exhbit decreased spreading and increased proliferation characteristic of the epithelial phenotype. Together, these results indicate that this parameter may serve as a suitable comparison to *in vitro* models of growth factor induced EMT. We thus compared these results to experiments in which EMT was induced by the soluble growth factor TGF-*β*1, as has previously been detailed [17]. Representative immunofluorescence images of MCF10A cells treated with increasing dosages of TGF-*β*1 illustrate a phenotypic switch from cortical actin, which is typically observed in epithelial cells, to pronounced actin stress fibers associated with the mesenchymal phenotype (Fig 4A). In these confluent monolayers, MCF10A average cell count decreases and average cell area increases for increase TGF-*β*1 doses (Fig 4B, D). As in Fig 3, we observe similar trends in simulations for decreasing cell contact inhibition (i.e., smaller *J*_*cc*_/*J*_*cm*_ ratio), although with a weaker dependence than observed *in vitro* (Fig 4C, E). Thus, we find that cell contact inhibition similarly regulates the cellular geometry averaged over the confluent monolayer in both simulation and experiment.

**Fig 4.**
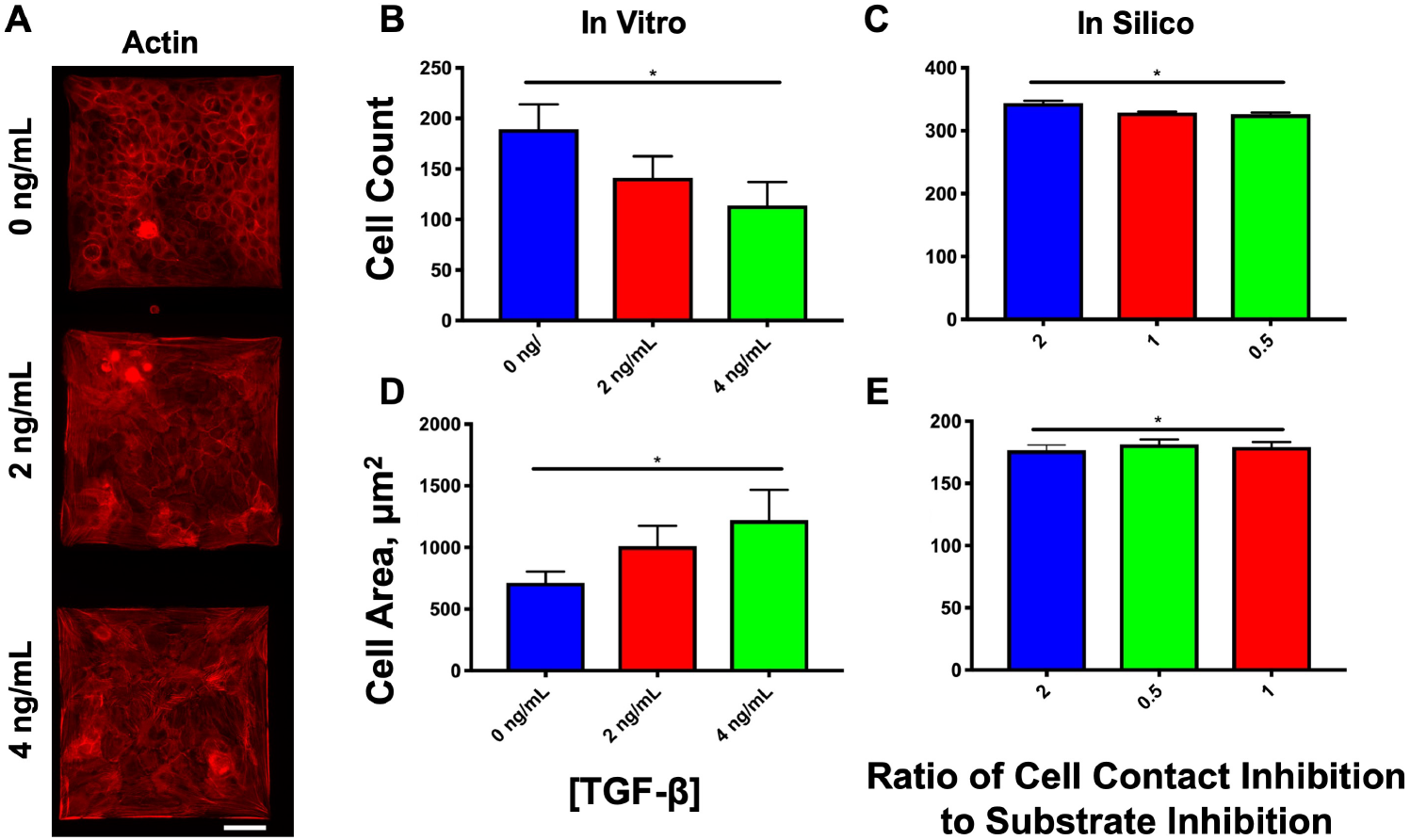
Morphological characterization of the epithelial phenotype with TGF-*β*1-induced EMT. (A) Representative immunofluorescent images of experimental illustrate a confluent MCF10A monolayer bounded to the 250 x 250 **µ*m* microfabricated square; scale bar = 50 **µ*m*. *In vitro* (B, D) and simulated (C, E) average cell count and cell area for the confined geometry are shown for each TGF-*β*1 dosage and ratio of contact interaction energies (*J*_*cc*_/*J*_*cm*_), respectively. Sample size *n*=3 for *in vitro* experiments. * with line denotes significance between each TGF-*β*1 dosage or each contact energy ratio.

### Cell-cell junction force maintains mechanical equilibrium of multicellular clusters

A key advance of the multicellular FMA model is the prediction of forces acting on cell-cell junctions. By assuming static equilibrium and applying a force-balance principle, cell-cell junction force was predicted as a reaction force that balances traction forces of the monolayer (described in detail in Methods). Cell-cell junction force magnitudes are shown on the boundaries between neighboring cells in simulated monolayers (Fig 5D). To examine spatial trends, we segmented the simulation domain into a 5 × 5 grid of bins and calculated the mean junction force magnitude within each bin (Fig 5E). The spatial distribution of junction forces is pronounced, with the largest forces in the interior and smallest in the corners (Fig 5F). However, interestingly, we find minimal variation in the spatial trends between low, medium, and high contact inhibition ratios.

**Fig 5.**
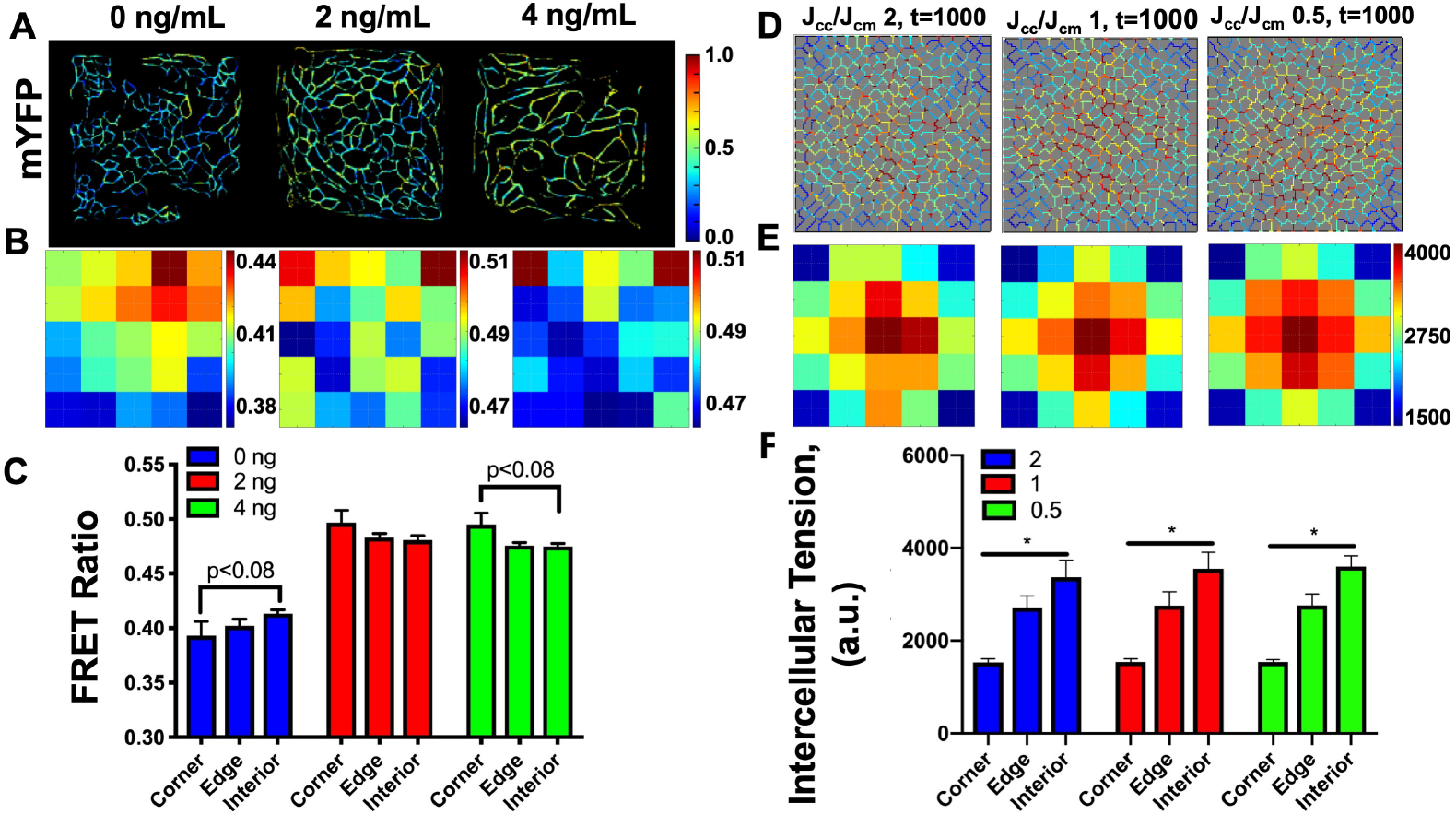
Intercellular interaction energy reflects TGF-*β*1 effects *in vitro*. (A) *In vitro* FRET intensities in MDCK II cells. (B) Corresponding heatmaps for average FRET intensities are binned into a 5 × 5 grid, and (C) their associated bar graphs averaged at the corners, edges, and interior for 0, 2, and 4 ng/mL TGF-*β*1 dosages; *n*=3. (D) Simulated intercellular tension is depicted as the net magnitude for high, medium, and low interaction energy (*J*_*cc*_/*J*_*cm*_) ratios. (E) Intercellular tension magnitudes are shown as a 5 × 5 grid with (F) their associated bar graphs averaged at the corners, edges, and interior; *n*=5, * with line denotes significance between each location.

We next sought to compare these with experimentally-measured junction forces. To measure cell-cell junction forces experimentally, Madin-Darby Canine Kidney Cells (MDCKII) cells were stably transfected with a full-length E-cadherin force sensor, as previously described [18]. Briefly, the force sensor consists of two fluorophores coupled by a polypeptide that exhibits elasticity. The two fluorphores are designed such that, when in close proximity, the pair exhibits Forster Resonance Energy Transfer (FRET): that is, emission light from the first fluorophore is absorbed by the second fluorophore, which emits light. As the sensor is stretched and the fluorophore pair moves apart, the excitiation of the second fluorophore by the first fluorophore decays, resulting in a loss of FRET excitation relative to excitation of the first fluorophore. This force sensor was inserted into E-Cadherin, which comprises the homophilic binding event in cell-cell junctions known as adherens junctions. Validation and functionality of this sensor has been previously demonstrated [19, 20]. EMT was again induced by increasing dosage of (TGF-*β*1) (Fig 5A). FRET ratio reflects the energy transfer between the two fluorophores, in which FRET ratio is inversely proportional to tension on the FRET force sensor: high FRET ratio indicates low tension and low FRET ratio indicates high tension. Representative pseudocolored images of the processed FRET ratio are shown in Fig 5A. We next investigated if spatial patterns of junction forces were established in these confluent monolayers. We again segmented images of the the local net FRET ratios into a 5 × 5 grid. In the absence of TGF-*β*1, colonies illustrated a nearly spatially uniform low FRET ratio, indicating high cell-cell tension throughout the monolayer (Fig 5B). TGF-*β*1 treatment increased FRET ratio, indicating a drop in overall tension. Additionally, a small spatial gradient was established, with higher FRET ratios (lower cell-cell tension) in the corner and edges and lower FRET ratios (higher cell-cell tension) in the interior of the monolayer, consistent with a spatial gradient of larger junction forces in the center and decreasing towards the edges and corners (Fig 5C).

Thus, we find that simulated cell-cell junction forces predict a spatial trend of decaying cell-cell tension from interior to periphery. Furthermore, simulated spatial gradients of cell-cell junction force are most comparable to experimental measures of TGF-*β*1-treated monolayers.

### Individual cell geometry spatial patterns

Summarizing our results presented thus far, we find that the multicellular FMA model reproduces contact inhibition-dependent trends for average cellular geometry (i.e., cell size and count), but underestimates this dependence compared with experimental observations. Further, our model qualitatively predicts trends for spatial patterns of cell-cell junction forces in TGF-*β*1-treated monolayers, but overestimates the magnitude of the spatial gradient, in comparison with experiments. We hypothesize that these discrepancies arise from an underestimation of cell size distribution throughout the monolayer in response to changes in contact inhibition. That is, individual cell size changes in response to TGF-*β*1 treatment due not only to loss of cell contact inhibition, but also to additional signaling not currently present in our model. To investigate this, we again segmented immunofluorescence images of MCF10A cells and binned cell area as before into a 5 × 5 grid (Fig 6A). Consistent with overall monolayer averages, cell area increased with increasing TGF-*β*1 dose. Evaluating the average cell area in the corner, edge, and interior of the monolayer reveals an overall increase in cell area at the periphery of the square, with the largest cell area localized to the corners in both low and high TGF-*β*1 dosages (Fig 6A). Reduced contact inhibition by treatment with TGF-*β*1 accentuates this trend, resulting in a large spatial gradient in cell area (Fig 6B).

**Fig 6.**
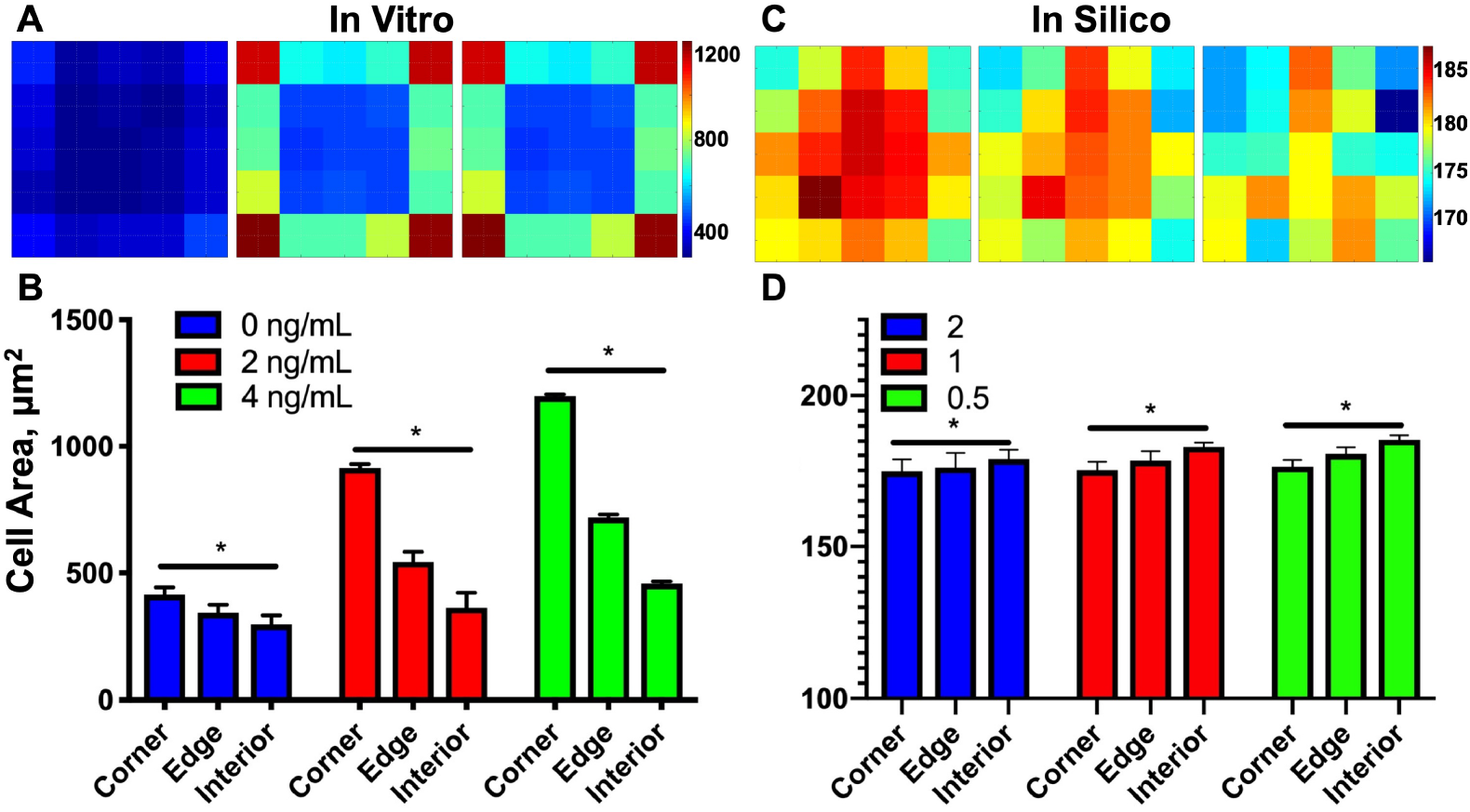
Individual cell geometry spatial patterns (A) *In vitro* heatmaps for binned cell area treated with 0, 2, and 4 ng/mL TGF-*β*1 and (B) their associated bar graphs for average corner, edge, and interior; *n*=3. (C) Simulated heatmaps for binned cell area at high, medium, and low contact inhibition and (D) their associated bar graphs; *n*=5.

In contrast, simulated cell area exhibited substantially reduced spatial variation compared to experimental cell area (Fig 6C). Furthermore, the effects of contact inhibition had a relatively minimal effect on spatial variation of cell area, resulting in slightly increased cell area at the monolayer interior (Fig 6D). Thus, the lack of accounting for heterogeneous cellular properties, specifically cell area, is a key limitation of our model. Since cells undergo profound phenotypic changes throughout EMT, it would be reasonable that these changes lead to parameter changes within the CPM for each individual cell; incorporating these changes in cell phenotype into the CPM component is a primary future goal for the model development.

### Analytical model of a simplified one-dimensional geometry

Both experimental and simulation data indicate that while traction forces are largest at the periphery of the epithelial cluster, junctional forces are largest near the center of the clusters and decay towards the periphery. We can gain additional insights by considering junction forces in tissue with a simple one-dimensional geometry, to both illustrate our approach and explain the perhaps counterintuitive prediction that larger traction forces at the periphery result in larger junction forces at the center. For this simple geometry, the traction and junction force magnitudes can be solved analytically, and further, these analytical results provide an explanation for some of the discrepancies between experiments and simulations noted above.

Consider a linear array of 2*n* cells of length *L* that are arranged and coupled in a line, such that the cell junctions are located at positions (−*nL*, 0), (−(*n*−1)*L*, 0),…, (0, 0),…, ((*n*−1)*L*, 0), (*nL*, 0), where *T* = *nL* is the length of half of the monolayer or tissue (Fig 7C). Note that the *y* position is insignificant, since all forces are oriented in the *x*-direction. The centroid of the tissue aligns with the origin, (0, 0), which is the junction on the left edge of cell 1, and thus the net traction force in each cell will be pointed towards this position. Further, we assume that each cell has *f* focal adhesions, uniformly spaced along the length of the cell *L*, and that traction forces are generated only at the focal adhesion positions. In the illustrated example, *f* = 4.

**Fig 7.**
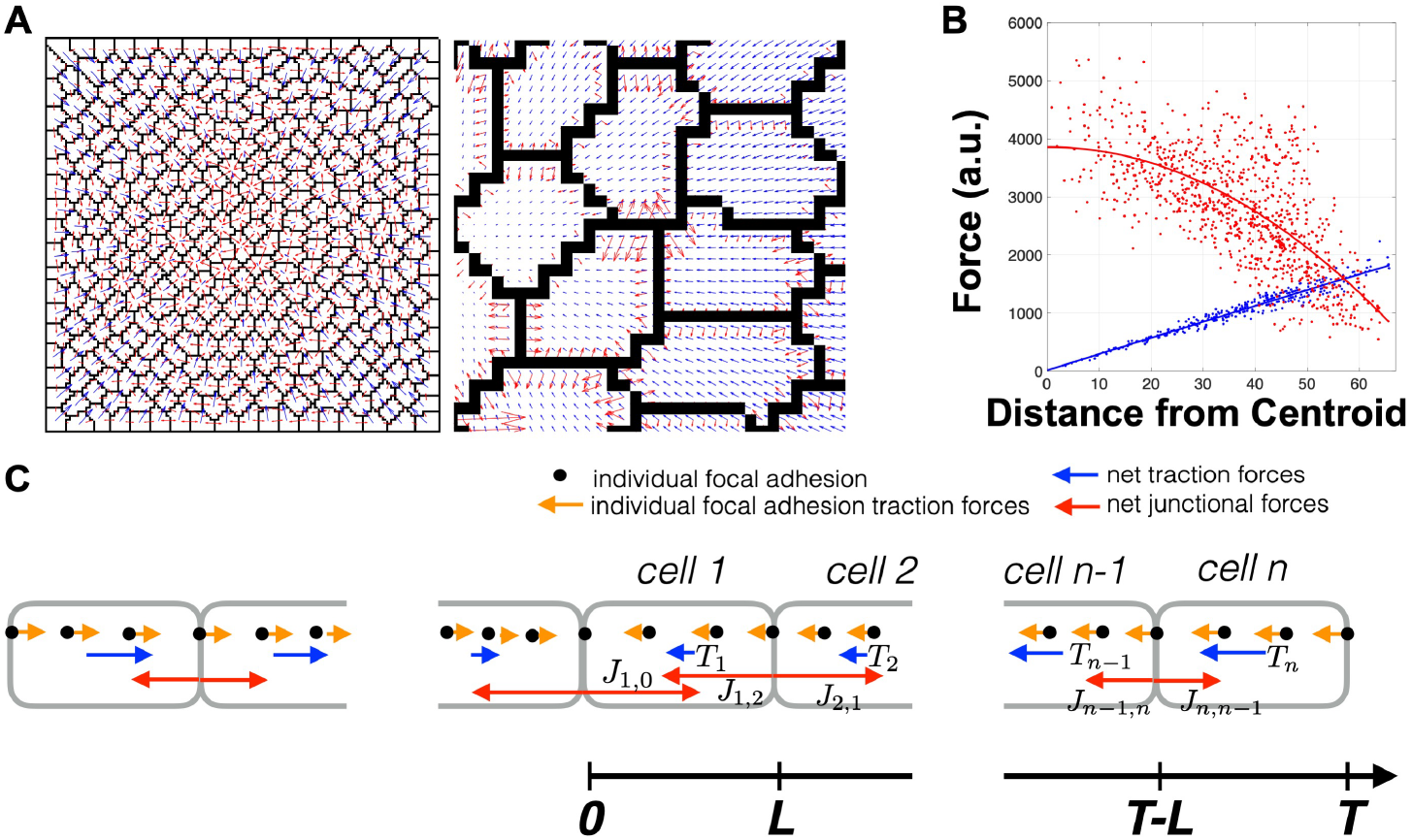
One-dimensional generalization for multicellular forces at mechanical equilibrium. (A) Representative snapshot of the traction and junction forces in the multicellular CPM model. (B) Plots of the traction and junction forces from the CPM simulations shows that traction force scales linearly with distance from monolayer centroid (blue line) and intercellular tension drops off quadratically from the centroid (red line).

Traction forces generated at each focal adhesion are thus proportional to distance from the origin, and the net traction force for a given cell is the sum of all traction forces over all focal adhesions. We can show that for cell *k*, with left edge at position ((*k* − 1)*L*, 0) and right edge at position (*kL*, 0), the net traction force is given by 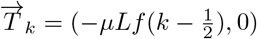, where *µ* is the appropriate scaling factor that relates cell geometry to traction forces [11]. For the rightmost cell, cell *n*, 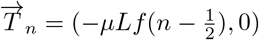. For mechanical equilibrium at cell n, this traction force must be balanced by the junction force from cell *n* − 1 to cell *n*, such that 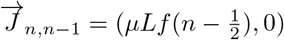. By assumption, net forces at the cell-cell junction are also in equilibrium, such that junction force pairs are symmetric, i.e., equal in magnitude and opposite in direction, such that 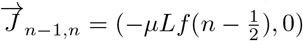.

Next considering forces on cell *n* − 1, the junction force from cell *n* − 2 to cell *n* − 1 must balance both the net traction force 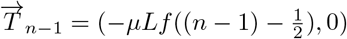 and junction force 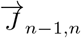, such that 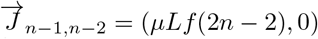. Similarly, junction force from cell *n* − 3 to cell *n* − 2, 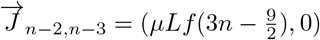. In general, we can show that the intercellular tension from cell *k* to *k* + 1,

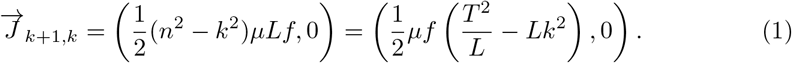

Thus, the junction force at the center onto the left edge of cell 1, 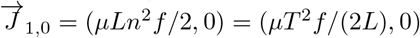. This simple geometry arrangement predicts larger magnitude junction forces in the center, and further illustrates a quadratic drop-off (due to the −*k*^2^ term in the magnitude of 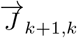) that is predicted as junction position k increases towards the periphery. A representative example of the CPM model illustrates the distribution of traction forces (blue) and junction forces (red) in a confluent monolayer (Fig 7B) and both the linear increase in traction force magnitude from the monolayer centroid and the quadratic drop-off in junction force magnitude (Fig 7B).

Thus, for a monolayer of a given size, i.e., fixed *T*, Eqn. 1 predicts that for a smaller cell size (decreased *L* and thus increased n), the magnitude of junction forces are larger throughout the monolayer, which is consistent with experimental measurements of lower FRET ratios (i.e., higher tension) in non-treated epithelial monolayers (Fig 5C). Further, in TGF-*β*1-treated monolayers, more mesenchymal-like larger cells at the monolayer periphery would be expected to have more focal adhesions per cell, in contrast with epithelial-like smaller cells in the interior. Additionally, while larger cells at the periphery will reduce junction forces locally, due to the cumulative nature of junction forces required to maintain mechanical equilibrium originating at the periphery, this local reduction in junction forces would be expected to have a greater influence on interior junction forces. All of these considerations would be predicted to reduce the magnitude of the spatial gradient, also consistent with smaller spatial gradients observed experimentally. Thus, we expect that our future work incorporating spatial variations in cell size in the CPM model will more accurately reproduce experimental results.

We can further generalize this example and consider the continuous limit in the spatial dimension, in which the traction forces *τ*(*x*) in the *x*-direction at position *x* (for *x* > 0) are given

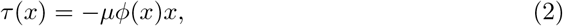

where *φ*(*x*) is the spatial distribution of focal adhesions per unit length. Junction forces *J*(*x*) at position *x* are then by definition the *second moment of area*, evaluated from the cluster periphery *T* to position *x*, where again *x* = 0 corresponds with the cluster center,

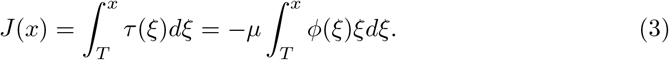

For uniform focal adhesion distribution, *φ*(*x*) = *f*/*L*, we can integrate Eqn. 2, and using the relationship *x* = *kL*, the result is equivalent to Eqn. 1.

## Discussion

In this study, we illustrate a generalized framework for predicting the spatial distribution of forces within and between cells in a monolayer. By assuming that i) clustered epithelial cells act as a syncytial unit and generate forces collectively in the FMA model; and ii) each cell in a monolayer exists in a quasi-equilibrium, in which junctional forces and traction forces are balanced, we are able to predict the distribution of cell-cell junction forces and cell traction forces within an epithelial cluster. Our model demonstrates that traction forces scale with the size of the multicellular cluster, a consequence of the FMA in which traction force is applied at uniformly distributed cell-matrix adhesions (i.e., at all pixels in the CPM). The model further predicts that the intercellular tension decays nonlinearly with the distance from the monolayer centroid. FRET analysis of mammary breast epithelial clusters indicate junction force distribution depends on monolayer geometry and not individual cell geometry, and confirms trends observed in simulations. The trends of our extended multicellular FMA model capture many key dynamical properties of monolayers undergoing EMT; however, the model does not capture spatial distribution trends observed in the control epithelial colony, possibly due to lack of consideration for heterogeneity of phenotype-specific cellular properties. The observed differences between simulations and experiments may owe to a number of factors, including nonuniformity in cellular phenotype that in turn alters cell size, and the number of cell-cell and cell-matrix attachments, as contacts between neighboring cells is not fixed.

A defining characteristic of TGF-*β*1-induced EMT is the disassembly of epithelial junctions, resulting in the loss of contact inhibition. During this process, intercellular tension redistributes from the cell-cell junctions to the cell-matrix attachments, which allows for increased mobility, growth, and spreading [15]. Our model represents this shift by altering contact penalties within the cell-cell and cell-matrix interaction energies. By altering the cell-cell contact energy, the model captures the contact inhibition of neighboring cells *in vitro*. However, simulating EMT via changes in the contact energy is not sufficient to capture all dynamics: in the CPM model, a defined value for optimal cell area constrains the simulated cell area that, in turn, limits cell-matrix adhesion. The shift from cell-cell contact to cell-matrix adhesion is indirectly restricted as a result. The spatial distribution of intercellular tension therefore predicts the spatial distribution of cell area, which would seem to indicate a shift towards cell-matrix adhesion. Future work will account for phenotype-dependent changes in the optimal cell area.

Many prior computational approaches have been developed to study tissue mechanical homeostasis and cell-matrix interactions. Vertex-based models, which consider mechanical force-balance along the boundaries of cells accounting for active and passive mechanical forces, have been developed to model tissue-scale emergent dynamics such as morphogenesis [21–23]. Agent-based models have been utilized to study cellular remodeling in response to mechanical perturbations, such as infarcts and wound healing [24–26]. The CPM framework has also been utilized to study cell-matrix interactions via extracellular matrix remodeling, in settings such as metastatic cancer cell migration and angiogenesis [27–29].

Our work builds on prior studies from Merks and colleagues that have demonstrated how local mechanical interactions can drive global cellular patterning and structure, using a hybrid CPM-FEM framework [10, 30, 31]. Multiscale modeling studies from Chaplain and colleagues have predicted that junction forces are redistributed as cells form colonies, which in turn can drive intracellular signaling pathways [32–34]. Interestingly, our extension to including multicellular mechanical interactions demonstrate that a gradient of intercellular tension can form even in the absence of heterogeneous cell populations. Through transduction of the mechanical gradient to intracellular signaling pathways, this tension distribution can provide positional information within a monolayer that regulates cellular phenomena, such as cell growth, proliferation, and migration. This is of particular interest to spatial regulation of EMT, during which cell stress is distributed to the monolayer periphery [35]. Connecting biochemical and mechanical signaling, the dependence on E-cadherin further suggests that intercellular tension may serve as a predictor of EMT.

Although the CPM predictions of force spatial distributions generally agree with previous findings, we find that model simulations do not fully capture monolayer dynamics observed *in vitro*. In particular, simulations do not reproduce spatial patterns in cell area. While TGF-*β*1 is known to increase cell spreading, the current model formulation defines a single target area for all cells, regardless of phenotype. As noted above, an ongoing focus of work is to incorporate variable cell target areas into the CPM to incorporate the effects of EMT on cell geometry and resulting spatial patterning in a more physiological manner.

## Materials and methods

In this study, we perform simulations and *in vitro* experiments to investigate intercellular tension and cell-matrix mechanical interactions in a multicellular geometry. Simulations were performed using a lattice-based cell model, the Cellular Potts model (CPM), generalized from the Potts model, to simulate epithelial monolayer dynamics [36]. The cell-occupied lattice is superimposed on a finite element lattice to determine substrate strains from simulated traction forces. In particular, we extend the first moment of area (FMA) prediction of single cell traction forces to predict the traction forces of a multicellular cluster. Lastly, we predict cell-cell junction forces by requiring that 1) cells in contact are mechanically coupled through cell-cell junctions, 2) the forces at these junctions balance net traction forces for each cell, and 3) the junction force is equal and opposite across a cell-cell adhesion. We compare model predictions of spatial patterning and junctional forces with *in vitro* experiments of TGF,*β*-treated epithelial cell monolayers.

### Cellular Potts model

The domain of the CPM lattice 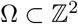 contains interconnected sites 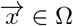 with spins 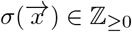 to identify the configuration of the domain. Each distinct cell-occupied site is defined by 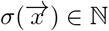, and an unoccupied site, i.e. extracellular matrix, is defined by 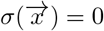. The CPM approximates the effective energy for a system configuration using a Hamiltonian term, where each term reflects a characteristic of biological cells and together summarize the configuration energy of the system. Here, the Hamiltonian is given by the sum of three terms

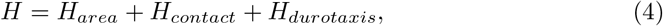

and Boltzmann statistics determine the probability of a possible lattice configuration

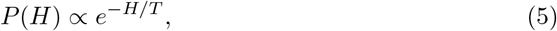

where *H* is the Hamiltonian defined in Eq. 4 and *T* > 0 is a temperature term that captures intrinsic cell motility.

The area term *H*_*area*_ approximates the cell area constraint as a deviation of the cell area relative to the target area such that

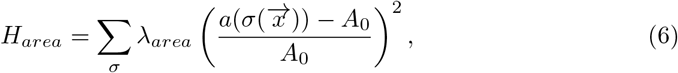

where 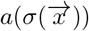 is the area of a given cell determined by number of lattice sites occupied by that cell, *A*_0_ = 312.50 *µ*m^2^ is the target area for all cells, and λ_*area*_ = 500 is an elasticity coefficient that maps deviations from the target area to a magnitude of energy.

The contact term *H*_*contact*_ represents costs due to contact between neighboring pixels, with different energies associated with cell-cell and cell-matrix interfaces:

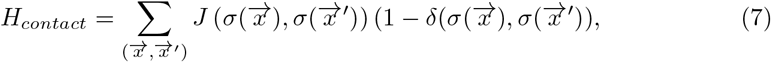

where 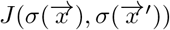, defines the interaction energy between adjacent lattice sites (*x*, *x′*) and 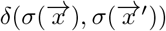 is the Kronecker delta function defined as 1 if a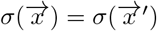 and 0 otherwise. We specify the cell-cell interface energy 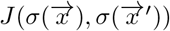 as *J*_*cc*_ and cell-matrix interface energy 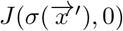 as *J*_*cm*_.

Lastly, the durotaxis term *H*_*durotaxis*_ introduced in van Oers [10] mimics the tendency for cell migration along gradients of mechanical strain. In particular, this term captures preferential cellular extension into lattice sites of higher strain

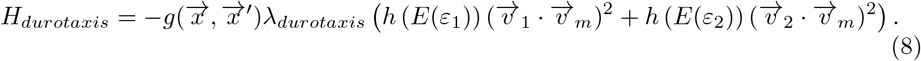

The *λ*_*durotaxis*_ = 1 term determines cell sensitivity to durotaxis; 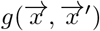 is 1 if a cell extends into a target site 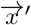 and −1 if a cell retracts; and the *v*_1/2_ ⋅ *v*_*m*_ are defined such that extension and retraction are greatest parallel to the major and minor principal strain axes, *v*_1_ and *v*_2_ respectively, and negligible perpendicular to it. The sigmoid function *h* (*E*) captures the preference for stiffer substrates,

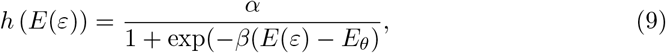

which assumes this preference has a minimal stiffness for spreading and reaches a maximum *α* = 10 at rate *β* = 5 × 10^−4^ kPa^−1^ and the half-max stiffness as E_*θ*_ = 15 × 10^3^ kPa. *E*(*ε*) is the cell perception of substrate strain stiffening,

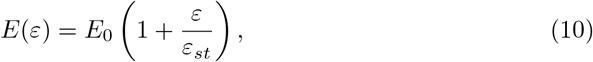

where *ε*_*st*_ = 0.1 determines the rate of strain-stiffening, *ε* is the substrate strain, and *E*_0_ = 10 kPa is the Young’s modulus of the substrate. The strain-stiffening only affects cell perception of strain-stiffening, not the stiffening of the finite element mesh itself (discussed below).

### Finite element analysis

To describe the substrate strain that governs durotaxis, we assume that a uniform, isotropic, and linearly elastic two-dimensional substrate deforms to cellular traction forces projected from the CPM (described below). The CPM lattice is mapped to the finite element lattice by relating each CPM lattice element to a finite element node. We solve the linear system

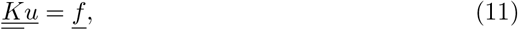

for the displacement *u* at each node, where *K* is the global stiffness matrix assembled from the stiffness matrix of each element, and *f* is the applied traction forces with constraint *u* = 0 at the CPM lattice boundary. In maintaining constant material properties during deformation, the element stiffness matrices 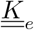 are given by

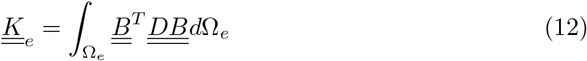

where *B* is the conventional strain-displacement matrix and *D* is the material property matrix under plane stress conditions

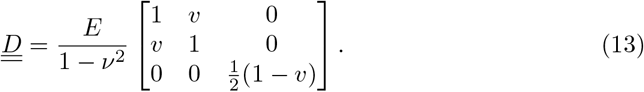

relating the Young’s modulus, *E* = 10 kPa, and Poisson’s ratio, *v* = 0.45, assuming planar stress. Lastly, *B* relates the local node displacements to the local strains by

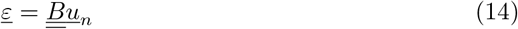

in which <u>*ε*</u> is a vector of the strain tensor 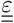.

### Traction forces

Prior work of van Oers and colleagues [10] assume that individual cell geometry relates to traction forces in the CPM by the first moment of area (FMA). Application of the FMA model to single cell geometries is previously described by one of the senior authors of this work [11]. In brief, the single cell FMA model assumes that each node *i* in a CPM cell *σ* exerts a force on all other nodes *j* in the same cell that is proportional to the distance between those nodes 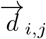,

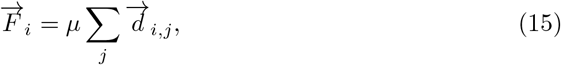

where *µ* is a scaling factor that relates cell geometry to traction forces. For simplicity, we assume *µ* = 1 nN *µ*m^−1^ and report forces as relative arbitrary units (a.u.). As shown in Lemmon and Romer [11], the resulting traction force at each CPM node is directed towards the cell centroid with magnitude proportional to the distance from the node to the centroid.

Here, we extend these previous works of the FMA model to describe the magnitude and direction of traction forces acting about a point in a multicellular geometry. For the multicellular FMA model, we assume that the boundary of two cells constitutes a cell-cell adhesion such that two or more adjacent cells behave as a single structural unit or cluster. We define an adjacency matrix *A*, where *A* is a *N*_*cell*_ × *N*_*cell*_ matrix, such that *A*_*σ*,*σ′*_ = 1 if cells *σ* and *σ′* are in contact, and 0 otherwise. By definition, *A* is symmetric. A cluster is defined as the connected components of the undirected graph defined by *A*.

Thus, the multicellular FMA model defines the traction force at each node in each CPM cell as directed towards the centroid of the associated multicellular cluster, with magnitude proportional to the distance from the node to the cluster centroid. Consistent with this hypothesis, recent experimental evidence supports an increase in traction forces with increasing multicellular cluster size [9, 37] For the case of a cluster comprised of a single cell, i.e., a cell lacking cell-cell adhesion, the multicellular FMA and single cell FMA model are equivalent.

### Intercellular tension

By construction, the single cell FMA model dictates that the sum of traction forces of an individual cell, i.e., the net traction forces 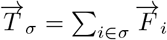 for cell *σ*, is equal to 0. In contrast, using the multicellular FMA model, the net traction forces of an individual cell *T*_*σ*_ within a cluster may not be equal to 0. Adapting a recent approach by Ng and colleagues [38], we hypothesize that junction forces are a reaction force, balancing the net traction force to maintain static equilibrium of each cell in a multicellular cluster. The multicellular FMA model is applied to calculate *T*_*σ*_ for each cell, and then we impose mechanical equilibrium on the multicellular clusters by relating the traction force to force across the cell-cell adhesion, such that for all cells *σ*,

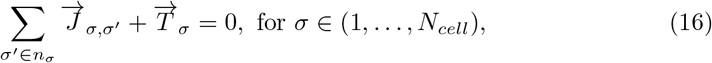

where *n*_*σ*_ defines the set of “neighbors” of cell *σ*, i.e., *A*_*σ*,*σ′*_ = 1, and *J*_*σ*,*σ′*_ is the junction force from cell *σ′* to cell *σ* (see S2 Fig). Eq. 16 defines *N*_*cell*_ linear equations, with 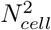 unknown *J*_*σ*, *σ′*_ terms. We further constrain the junction force calculations by assuming that junction force pairs are equal in magnitude and opposite in direction, i.e.,

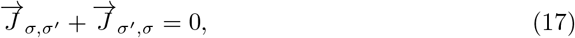

for all (*σ*, *σ′*) such that *A*(*σ*, *σ′*) = 1.

Combining Eqs. 16 and 17, we arrive at a linear system with a set of *N*_*cell*_ + *N*_*junc*_ equations and 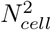 unknowns (see S2 Fig), where *N*_*junc*_ is the number of intercellular junctions, which can be determined by the sum of the terms above (or below) the main diagonal of *A*, with a maximum value of *N*_*cell*_(*N*_*cell*_ − 1)/2. In practice, linear systems for Eqs. 16 and 17 are determined separately to both the x- and y-components of the traction and junction forces.

For nearly all cluster arrangements, the resulting linear system is overdetermined. Analogous to the CPM thermodynamic energy minimization, we assume that the solution to be the minimization of junction force for each cell pair in the cluster, such that *J*_*σ*, *σ′*_ terms are calculated as the minimum norm least-squares solution to the linear system (using the MATLAB lsqminnorm function).

### Cell division

We incorporate cell division into the CPM model to reproduce epithelial cell capacity to proliferate and form a confluent monolayer. For simplicity, we assume that if an individual cell area exceeds a minimum area threshold, which we define as 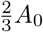, then individual cells divide with random probability *p*_*divide*_ = 0.005, unless otherwise stated. For cell division, following the prior approach of Daub and Merks, we compute the line of division for each CPM cell as the line following the minor axis, such that each daughter cell is of approximately equally area [28].

### Numerical simulations

The CPM map is initialized as uniformly distributed pixels of size 100 × 100, for which each pixel corresponds with a size of Δ*x* = 2.5 *µ*m. Initial seeding is dispersed on the cell map excluding the outermost boundary with random probability, *p* = Δ*x*/(4*A*_0_). An unloaded finite element mesh of size 101 × 101 forms the nodes of attachment for cells of the CPM map, in which each cell-occupied pixel occupies four nodes. To calculate forces from the CPM map, pixels are first mapped to the finite element substrate by identifying the corresponding nodes. At a given instant, the single cell or multicellular geometry is sufficient to define cellular traction forces at each node, using the single or multicellular FMA model, as described above, respectively. The resulting traction forces govern the displacement at each node and determines the strain in the finite element mesh, which in turn is used in evaluating *H*_*durotaxis*_.

Cell movement consists of copy attempts of randomly selected pixel at each Monte Carlo step (MCS). For each pixel to have equal probability of selection, each MCS has a total of 10^4^ copy attempts. For each copy attempt, a pixel is selected and randomly perturbed; the sum of interaction energies with each pixel in the Moore neighborhood, Σ*J*(*σ*(*x*, *x′*)), determines the *H*_*contact*_ term. Lastly, the cell area before and after the copy attempt provides the *H*_*area*_ term. Together, the net change in the Hamiltonian associated with that copy attempt, i.e. Δ*H*(*σ*(*x*, *x′*)), provides the local energy for the cell before and after the copy attempt. The copy attempt is accepted (*σ*(*x*) → *σ*(*x′*)) with probability determined by the partition function (Eq. 5) for Δ*H* > 0 and probability 1 for Δ*H* < 0.

For parameter analysis, the parameter set consisted of each combination of cell-cell interaction energies and cell-matrix interaction energies, *J*_*cc*_ and *J*_*cm*_, respectively, each repeated with a uniquely seeded random number. The confluence is determined by the ratio of total cell occupied pixels to the total grid area. The cell area is number of pixels occupied by each unique cell state, and the cell count is the number of unique states.

### Cells and reagents

All cells were cultured in a humidified atmosphere at 37°C with 5% CO_2_. Human MCF10A mammary epithelial cells were obtained from the National Cancer Institute Physical Sciences in Oncology Bioresource Core Facility, in conjunction with American Type Culture Collection (Manassas, VA). MDCK II cells were a gift of Rob Tombes (VCU). MCF10As were maintained under standard culture conditions in DMEM/F-12 HEPES (Life Technologies, Carlsbad, CA), supplemented with 5% horse serum, 0.05% hydrocortisone, 0.01% cholera toxin, 0.1% insulin, 0.02% EGF and 1% antibiotics. MDCK II cells were maintained under standard culture conditions in DMEM (Life Technologies, Carlsbad, CA) supplemented with 10% fetal bovine serum and 1% antibiotics. Purified recombinant active TGF-*β*1 was purchased from Sigma Aldrich (St. Louis, MO). Immunofluorescence imaging was conducted using the following primary antibodies: Ms anti-Hu E-cadherin (HECD-1, Abcam, Cambridge, United Kingdom), Ms anti-Ms N-cadherin (BD Biosciences, San Jose, CA), Rb anti-Hu FN (Abcam, Cambridge, United Kingdom), Ms anti-Hu LTBP-1 (RD Systems, Minneapolis, MN), Rb anti-Hu Smad2 (86F7, Cell Signaling Technology, Danvers, MA), Dapi (Thermo Fisher Scientific, Waltham, MA). F-actin images were acquired by labeling cells with AlexaFluor555 Phalloidin (Life Technologies, Carlsbad, CA).

### Microcontact printing

Microcontact printed square islands were generated as previously described [39]. Briefly, 250 *µ*m × 250 *µ*m squares were constructed by generating a negative mold template on a silicon wafer made from an epoxy-type, near-UV photoresist (SU-8; Microchem) using traditional photolithographic techniques. A replica-mold of poly(dimethylsiloxane) (PDMS; Sylgard 184, Fisher Scientific, Hampton, NH) raised patterns were be coated with 100 µg/ml laminin (Sigma Aldrich, St. Louis, MO) for 2 hours at 37 degree C. Stamps were then rinsed in dH2O and dried with nitrogen gas. The laminin square islands were then stamped onto a thin layer of UV-treated PDMS on top of a glass coverslip. 2% Pluronics F-127 in phosphate-buffered saline (PBS) was used to prevent cells from adhering outside of the laminin-stamped areas. Coverslips were rinsed in PBS prior to cell seeding. Efficiency of protein transfer was confirmed by Immunofluorescence labeling of the ECM protein.

### Immunofluorescence microscopy

MCF10A and MDCKII cells were plated on microcontact-printed laminin islands at cell densities that resulted in near-confluent monolayers. After 6 hours, samples were rinsed in culture medium to remove non-adherent cells. Cells were cultured for 18 hr and were then transferred to EGF- and serum-free culture conditions for 2 hr to induce an epithelial phenotype. Cells were then incubated with or without TGF-*β*1 for an additional 48 hours. Cells were permeabilized with 0.5% Triton in 4% paraformaldehyde for 2 minutes, then incubated in 4% paraformaldehyde for 20 minutes. Several PBS-rinses were performed, followed by blocking in 0.1% BSA and labeling with primary antibody for 30 minutes at 37 degree C. Cells were then blocked again in 0.1% BSA and incubated with the appropriate secondary antibody for 30 minutes. Images were acquired on a Zeiss AxioObserver Z1 fluorescence microscope using ZEN2011 software.

### Cell area and cell number quantification

Cell area and cell number were determined by analyzing immunofluorescence images of F-actin and nuclei via an custom-written image processing algorithm in MATLAB.

Binary masks of nuclei were generated by thresholding grayscale nucleus images; objects in the binary mask were counted to determine total cell number. To determine cell size, the centroid of each object in the binary mask was determined using the regionprops function. Nuclei centroids were used to generate a Voronoi diagram, which consists of a series of polygons that have edges that are equidistant from neighboring nuclei.

Previous studies have demonstrated that Voronoi diagrams reasonably predict cell boundaries in an epithelial monolayer [40], and provide a more consistent quantification of cellular size as opposed to quantification of protein markers in the cell-cell junction, whose expression and localization changes as TGF-β dose increases. Cell area was calculated for each cell by summing the pixels in each Voronoi polygon, and were averaged across the 250 *µ*m × 250 *µ*m colony. Spatial localization of cell number and cell area were determined by binning nucleus centroids into a 5 × 5 grid. Cell counts in each bin were totaled, and cell areas for each bin were averaged if the nuclei centroid was contained within the bin. Spatial localization data was further combined into either corner bins, edge bins, or interior bins, such that there were no overlap between the three regions (i.e., corner bins were not included in the edge region).

### FRET analysis

To measure force on cell-cell junctions, Fluorescence Resonance Energy Transfer (FRET)-based, full-length E-cadherin tension biosensors were stably transfected into MDCK II cells. Epithelial square islands were cultured as stated above, and images were acquired on a Zeiss LSM 710 laser scanning microscope using ZEN2011 software. Briefly, mTFP (donor) and mEYFP (acceptor) fluorophores were imaged utilizing spectral unmixing at 458 nm excitation. The acquired intensity images were manually masked through ImageJ. Background subtraction and removal of saturated pixels was then performed via an image processing algorithm in Python as previously described [41]. FRET ratio was determined by obtaining the acceptor/donor ratio and multiplying with a binary mask of the junctions. This allowed for inspection of FRET pixels of interest within outlined cell-cell junctions.

### Statistical analysis

Simulated and experimental data was exported to Prism 8 (GraphPad Software Inc) for analysis. Statistical significance, indicated by a p-value less than 0.05, was determined by one-way ANOVA across each TGF-*β*1 dosage, ratio of interaction energies, and/or spatial localization.

## Supporting information

**S1 Fig. Spatial maps of substrate strain.** Spatial maps of substrain strain are shown for simulated cells in which traction forces are calculated based on (A) individual cell geometries or (B) multicellular clusters. Maps correspond with simulations shown in Fig. 1.

**S2 Fig. Colony connectivity and intercellular tension.** Simplified depiction of four neighboring cells (gray) forming a multicellular cluster and the corresponding adjacency matrix, A (left). Traction forces (red arrows) are proportional to the FMA about the centroid of the cluster (green dot). Junction forces (blue arrows) balance the net force imbalance for a given cell. The linear system is constructed from the mechanical equilibrium matrix and junction symmetry matrix (right). The mechanical equilibrium matrix is constructed from the connectivity of each cell given by the adjacency matrix and by applying the force balancing principle. The junction symmetry matrix requires each junction force across a cell-cell adhesion to be equal and opposite.

**S1 Video. Single cell without proliferation.** Simulated cell organization for the single cell FMA model as shown in Figure 1A. Movie corresponds to simulation of 1000 Monte Carlo Steps.

**S2 Video. Multicellular without proliferation.** Simulated cell organization for the multicellular FMA model as shown in Figure 1B. Movie corresponds to simulation of 1000 Monte Carlo Steps.

**S3 Video. In vitro proliferation.** Spatiotemporal dynamics of MCF10A cells confined to a 250 *µ*m × 250 *µ*m PDMS square as shown in Figure 2A. Movie corresponds with experiments of 24 hours.

**S4 Video. Multicellular CPM with proliferation.** Spatiotemporal dynamics of simulated cells for the multicellular FMA model with cell division probability of 0.5% per time step as shown in Figure 2B. Movie corresponds to simulation of 1000 Monte Carlo Steps.

**S1 Table. Model parameters.** Key model parameters and values are shown. * Value used unless otherwise noted.

